# PTEN loss in glioma cell lines leads to increased extracellular vesicles biogenesis and PD-L1 cargo in a PI3K-dependent manner

**DOI:** 10.1101/2023.07.26.550575

**Authors:** Julio C. Sanchez, Timothy M. Pierpont, Dariana Argueta-Zamora, Kristin Wilson, Avery August, Richard A. Cerione

## Abstract

Phosphatase and Tensin Homologue (PTEN) is one of the most frequently lost tumor suppressors in cancer and the predominant negative regulator of the PI3K/AKT signaling axis. A growing body of evidence has highlighted the loss of PTEN with immuno-modulatory functions including the upregulation of the programmed death ligand-1 (PD-L1), an altered tumor derived secretome that drives an immunosuppressive tumor immune microenvironment (TIME), and resistance to certain immunotherapies. Given their roles in immunosuppression and tumor growth, we examined whether the loss of PTEN would impact the biogenesis, cargo, and function of extracellular vesicles (EVs) in the context of the anti-tumor associated cytokine interferon-γ (IFN-γ). Through genetic and pharmacological approaches, we show that PD-L1 expression is regulated by JAK/STAT signaling, not PI3K signaling. Instead, we observe that PTEN loss positively upregulates cell surface levels of PD-L1 and enhances the biogenesis of EVs enriched with PD-L1 in a PI3K-dependent manner. We demonstrate that because of these changes, EVs derived from glioma cells lacking PTEN have a greater ability to suppress T cell receptor (TCR) signaling. Taken together, these findings provide important new insights into how the loss of PTEN can contribute to an immunosuppressive TIME, facilitate immune evasion, and highlight a novel role for PI3K signaling in the regulation of EV biogenesis and the cargo they contain.

## Introduction

Glioblastoma multiforme (GBM) is classified as a grade IV astrocytoma by the World Health Organization and is the most common and aggressive CNS-related tumor with a mean survival period ranging between 12-18 months.^1^ With a standard of care that has not changed in over two decades, and only 6-12% of patients reaching the five-year survival mark, drastically underscores a need to bring new strategies to bear on this devastating disease.^2^ The advent of novel immunotherapies such as immune checkpoint inhibitors (ICIs) has shown a great deal of success against certain forms of cancer by successfully treating otherwise terminally ill patients.^3^ ICIs are monoclonal antibodies that block the interaction between immunosuppressive ligands, typically expressed by tumor or myeloid cells, and their cognate checkpoint receptors expressed on T cells. For example, blocking the interaction between the programmed death protein 1 (PD-1) receptor and its ligand (PD-L1) allows for adequate T cell receptor (TCR) activation and thus anti-tumor cytotoxic T cell functions to manifest^3^. For this reason, studies elucidating how PD-L1 and other immunosuppressive ligands are regulated at the cellular level have garnered a great deal of attention. Cancer cells upregulate PD-L1 expression in response to inflammatory cytokines such as interferon-γ (IFN-γ), in a process known as “adaptive immune resistance,” and can be predictive of successful patient response rates to ICI.^4,5^ However, while ICIs have led to robust response rates in select tumor types such as melanoma and NSCLC, the majority of patients bearing solid tumors, including those diagnosed with GBM, have experienced little benefit from these therapies.^6–8^ For GBM, several factors influence the effectiveness of ICI and other immunotherapies including low tumor mutational burden, few tumor infiltrating lymphocytes (TILs), a significantly large population of immunosuppressive tumor associated macrophages (TAMs), and their utilization of various methods for immune evasion.^9–12^ Recent clinical reports have associated the loss of PTEN as a significant biomarker for patients who fail to respond to ICIs across several tumor types including GBM, melanoma, gastrointestinal and breast.^13–16^ PTEN is one of the most frequently mutated and/or deleted tumor suppressors across all cancers,^17,18^ with 40-60% of GBM patients exhibiting such deletions.^19^ Therefore, understanding how the loss of PTEN impacts the tumor immune microenvironment (TIME), immune evasion, and its contribution to ICI resistance, represent critically important questions within the field.

PTEN is recognized as the primary lipid phosphatase responsible for reverting the class I PI3K- generated lipid second messenger, phosphatidylinositol (3,4,5)-triphosphate (PIP3), to phosphatidylinositol (4,5)-diphosphate (PIP2).^20^ The loss of PTEN and/or hyperactivating mutations associated with PI3K lead to elevated levels of PIP3 and increased activation of important pleckstrin homology (PH) containing protein kinases such as PDK1 and AKT, as well as disabling the negative regulation of mTORC2 by mSin1. Together, these protein kinases activate many of the oncogenic signaling pathways that give rise to enhanced cell growth, survival, metabolic reprogramming, and cell migration.^21–23^ While loss of PTEN is most recognized for promoting the signaling activities downstream of PI3K, a growing body of evidence suggests that impaired PTEN function and the accompanying enhancement in the actions of PI3K activity contribute toward the development of a more immunosuppressive TIME. Indeed, tumors lacking PTEN have been associated with an immunologically “cold” TIME, harboring fewer TILs and more immunosuppressive and pro-tumorigenic myeloid and lymphoid cells which can negatively impact immunotherapy treatment outcomes. ^24–26^ These observations suggest that the loss of PTEN may influence how tumor cells communicate with their immediate environment. Indeed, it has been observed that loss of function associated with PTEN can impact the secretome of glioma and other cancer cells, including chemokines and cytokines that impact tumor growth, metastasis and even the intravasation and reprogramming of lymphoid and myeloid cells.^27–31^ These findings thus prompted the question of whether the loss of PTEN and/or hyperactive PI3K signaling might also affect the biogenesis and/or cargo of extracellular vesicles (EVs), which represent another major component of the cancer cell secretome.

EVs are a group of secretory vesicles that play important roles in intercellular communication in a wide range of biology and disease through receptor-ligand interactions and as a result of the horizontal transfer of protein and nucleic acid cargo to the cells they engage.^32,33^ There are two major classes of EVs, designated as microvesicles or large EVs (ranging from 250 nm to 1 micron in diameter) which mature and bud off from the cell surface, and exosomes or small EVs (typically less than 200 nm in size) that form as intraluminal vesicles within multi-vesicular bodies (MVBs) and are released from cells upon the fusion of MVBs with the plasma membrane.^34,35^ EVs have been extensively studied in cancer and shown to play important roles in shaping the TME by transforming surrounding stromal cells, and through their ability to induce angiogenesis, establish a pre-metastatic niche, and help mediate immune evasion.^36–39^ Indeed, EVs derived from both GBM and melanoma cells have been shown to locally and systemically suppress the activation of CD8^+^ T cells in a PD-L1-dependent manner^40,41^ and to alter the fate of TAMs.^42^ Interestingly, several reports using different cancer cell types, including GBM, have implicated hyperactive PI3K signaling and/or PTEN loss with an increase in PD-L1 expression at the cellular level.^43–47^ Therefore, we were interested in investigating how PTEN/PI3K signaling might influence the biogenesis of EVs and the presence of PD-L1 as EV-associated cargo.

Using four isogenic glioma cell lines, we demonstrate that following exposure to IFN-γ, cells lacking PTEN have a marked increase in both surface levels of PD-L1 and as EV associated cargo. We further show that in the absence of PTEN, cells exhibit an increase in total EV biogenesis. Both the restoration of PTEN and the pharmacological inhibition of PI3K reversed these effects, indicating that EV biogenesis and PD-L1 cargo sorting are regulated in a PI3K-dependent manner. Using a Jurkat reporter cell line, we then demonstrate that EVs shed from glioma cells are capable of inhibiting TCR signaling in a dose-dependent manner with this effect being most pronounced when vesicles are shed from cells that lack PTEN.

## Results

### PTEN impacts cell surface levels of PD-L1 but not global protein expression

PTEN loss in GBM has been associated with increased expression of PD-L1 in steady-state conditions as a result of enhanced PI3K/AKT activity.^43^ However, given the recent appreciation that IFN-γ has a role in upregulating PD-L1 and in treatment outcomes relating to ICI, we re-examined the relationship between PTEN status and PD-L1 expression in four glioma cell lines (U87, U251, LN229, and T98G) with previously confirmed PTEN status.^43,48^ Using lentiviral transduction, we induced the ectopic expression of PTEN in U251 and U87 cells to determine if differences in PD-L1 expression existed with or without IFN-γ treatment. Twenty-four hours post-treatment, cells were lysed and subjected to immunoblot analysis. As expected, PTEN expression reduced PI3K activity as indicated by decreased levels of phosphorylated AKT (p-AKT) (Figure 1A). Additionally, although there was a clear and marked upregulation of PD-L1 expression in response to IFN-γ treatment, there was no clear indication that PTEN/PI3K signaling impacted overall PD-L1 expression levels regardless of IFN-γ treatment (Figure 1A). To determine whether the same was true at the transcriptional level, we performed qPCR analysis on the isogenic U87 and U251 cells with or without IFN-γ. While IFN-γ treatment gave rise to an expected increase in PD-L1 transcripts, no difference in PD-L1 transcription was observed in a PTEN-dependent manner, regardless of IFN-γ treatment (Figure S1A). To broaden our comparison, we used CRISPR technology to knock out (KO) *PTEN* in LN229 and T98G cell lines across. Apart from the T98G cell line, which harbors a point mutation in *PTEN* that impairs its function and thus increases AKT activity, ^48^ the manipulation of PTEN across the three other cell lines corresponded with the expected changes in the phosphorylation status of AKT (Figure 1B). The ectopic expression of PTEN in U87 and U251 cells resulted in reduced p-AKT levels, whereas the KO of *PTEN* in LN229 cells led to increased p-AKT. Importantly, across all cell lines, PTEN/PI3K signaling did not appear to affect PD-L1 expression in response to IFN-γ (Figure 1B). Because IFN-γ can be present in acute or chronic contexts within a tumor, we examined whether temporal differences in IFN-γ exposure would lead to divergent PD-L1 expression patterns in U87 cells in a PTEN-dependent manner; however, no obvious differences in PD-L1 levels were observed (Figure S1B). Taken together, these data suggested that the upregulation of PD-L1 in response to IFN-γ was occurring in a PTEN-independent manner.

**Figure 1.**
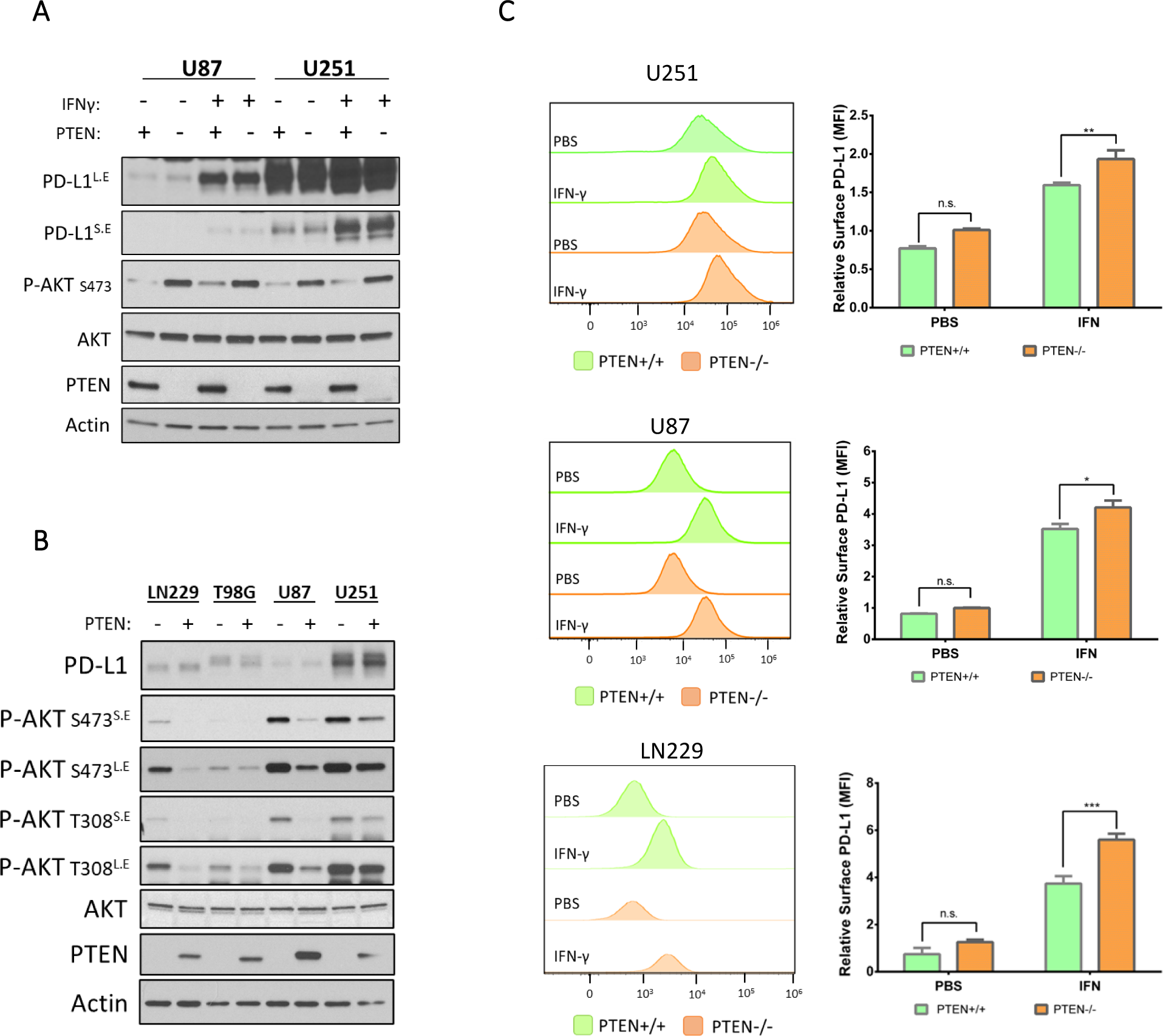
PTEN loss leads to increased cell surface levels of PD-L1 after IFN-γ exposure. A) Semi-stable U251 and U87 cells transduced with empty vector control or with PTEN vector were treated once with IFN-γ (50 ng/mL) or with PBS as a control. Twenty-four hours later, cells were serum starved for 12-15 hours, lysed, and subjected to immunoblot analysis to compare expression of PD-L1, PTEN, total AKT, and p-AKT (S473). Actin was used as a loading control. Some blots are shown as long exposure (L.E) and short exposure (S.E) to improve dynamic range. B) Semi-stable isogenic glioma cell lines were created by either CRISPR-mediated PTEN KO (LN229 and T98G cells) or by the ectopic expression of PTEN (U87 and U251 cells). All cells were treated once with IFN-γ (50 ng/mL). After 24 hours, cells were serum starved for 12-15 hours, lysed, and subjected to immunoblot analysis to compare the relative expression of PD-L1, PTEN, and AKT activation (T308, S473), with actin serving as a loading control. Long exposure (L.E), short exposure (S.E). C) U251, U87 and LN229 were treated with IFN-γ (50 ng/mL) or PBS control for 24 hours to evaluate changes in cell surface PD-L1. Cells were treated with αPD-L1-PE antibody or isotype control antibody and analyzed by flow cytometry. Left, representative histograms for each cell line showing PD-L1 expression on the X-axis represented by fluorescence intensity relative to the number of cells analyzed on the Y-axis. Right, PD-L1 expression is represented by the mean fluorescence intensity (MFI). Values from 3 different experiments were averaged and analyzed for statistical significance. Data for LN229 cells were acquired from four clones, two with PTEN expression and two without. Statistical analysis for all graphs was done using 2-way ANOVA with Tukey multiple comparisons test. *p-value 0.0190, **p-value 0.0097, *** p-value 0.001

It is known that IFN-γ binds to its cognate receptor to stimulate the JAK/STAT class of transcription factors to upregulate PD-L1 expression.^49^ Therefore, we concomitantly treated U87 cells with IFN-γ, tofacitinib (a JAK/STAT inhibitor), or LY294002 (a PI3K inhibitor), to determine which signaling pathway was necessary for PD-L1 expression. Using immunoblot analysis, we again determined that reduced PI3K activity failed to inhibit PD-L1 expression regardless of IFN-γ treatment (Figure S1C). In contrast, tofacitinib successfully blocked IFN-γ mediated expression of PD-L1 in a dose-dependent manner (Figure S1D). Collectively, these data further indicated PD-L1 expression was independent of PI3K/AKT signaling and that IFN-γ mediated PD-L1 expression is dependent upon JAK/STAT signaling.

Although we had not observed differences in overall PD-L1 expression in a PTEN-dependent manner, we wanted to evaluate if any changes in PD-L1 were occurring specifically at the plasma membrane (PM). Using flow cytometry, we analyzed PD-L1 levels on three isogenic cell lines for *PTEN*, (LN229, U87, and U251 cells). Twenty-four hours after IFN-γ treatment all cells showed an increase in cell surface PD-L1 expression, however, cells expressing PTEN showed a marked decrease in PD-L1 compared to their PTEN-null counterparts (Figure 1C). The fact that PTEN appeared to negatively influence the amount of PD-L1 present at the PM of cells raised the possibility that PTEN may also be playing a role in the endocytic trafficking or regulation of PD-L1 directed toward the PM. Indeed, it is beginning to be appreciated that PTEN can regulate endosomal maturation to influence and regulate mitotic signaling.^50^ Because EVs originate from either the PM or endomembrane compartments, this led us to investigate whether the loss of *PTEN* and/or reduced PI3K signaling could influence EV biogenesis and the inclusion of PD-L1 as cargo.

### PI3K inhibition and PTEN expression negatively impact extracellular vesicle biogenesis

We next explored whether PI3K/PTEN signaling and/or IFN-γ could impact EV biogenesis. Using the isogenic U87 cell line for PTEN expression and a pharmacological PI3K inhibitor (LY294002), we confirmed through immunoblotting that both PTEN expression and LY294002 led to a corresponding decrease in p-AKT, while IFN-γ treatment promoted the upregulation of PD-L1 expression (Figure 2A). The conditioned media from these cells was processed by differential centrifugation (Figure S2A) to generate partially clarified media (PCM) which was subjected to nanoparticle tracking analysis (NTA) to determine whether the overall diameter and/or relative abundance of EVs changed in response to IFN-γ and/or PTEN status. Notably, we observed a 50% reduction in EV biogenesis in response to PI3K inhibition from both pharmacological and genetic approaches (Figure 2B, upper panel). In contrast, IFN-γ treatment did not have a significant effect on the numbers of EVs shed from U87 cells regardless of their PTEN status (Figure 2B, lower panel).

**Figure 2.**
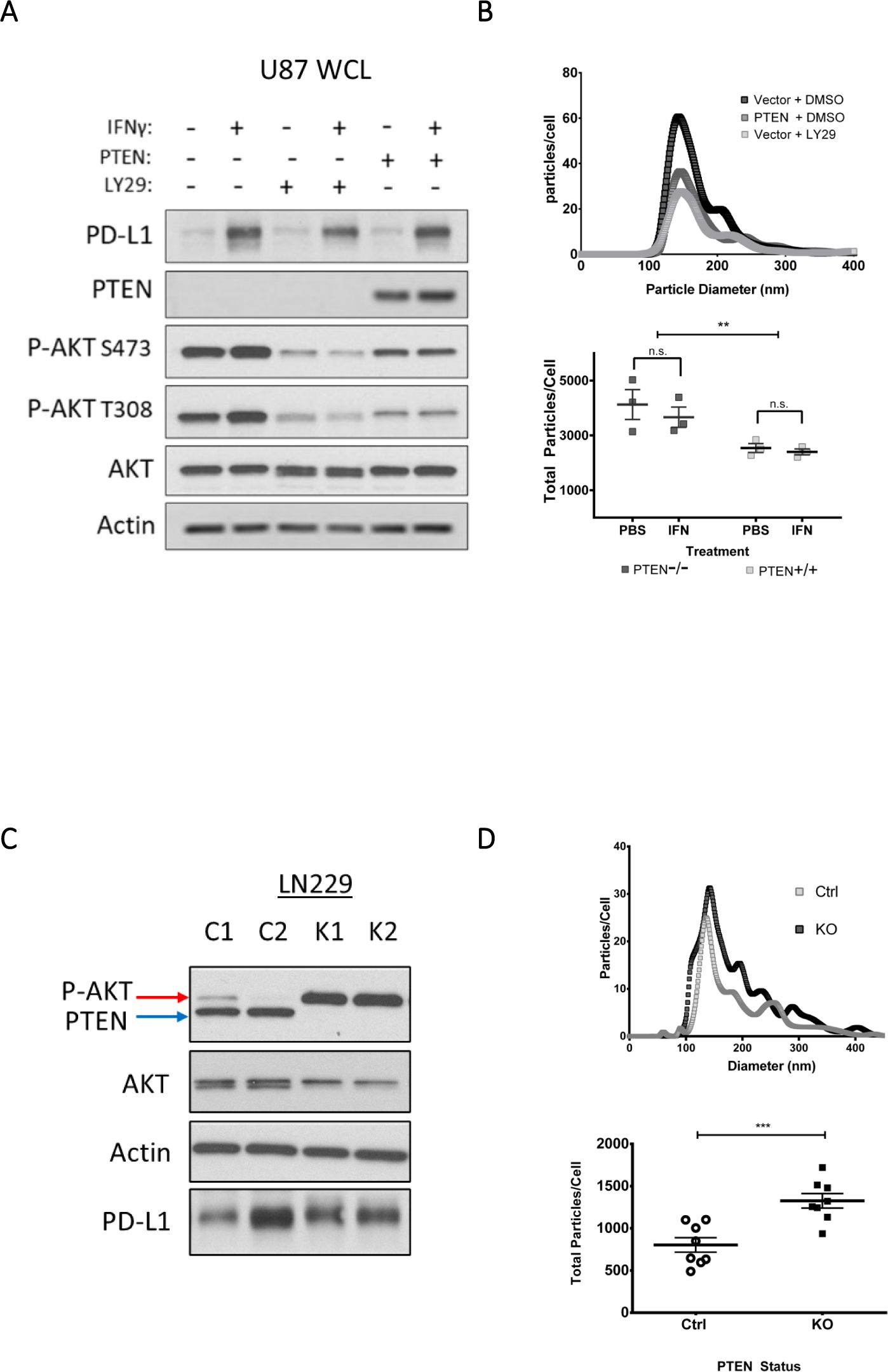
Elevated PI3K signaling as a result of PTEN loss is associated with increased EV biogenesis in isogenic LN229 and U87 cell lines. Semi-stable U87 cells were concomitantly treated with the PI3K inhibitor, LY294002 (LY29, 30 μg/mL) or with DMSO as a vehicle control and with IFN-γ (50 ng/mL) or PBS as the vehicle control. Twenty-four hours post-treatment, cells were washed three times and cultured for 12-15 hours in serum-free base media. The conditioned media was collected for downstream analysis while cells were lysed and analyzed by immunoblot to compare PTEN, total AKT, phosphorylated AKT (T308, S473), and actin as a loading control. The conditioned media from “A” was collected and processed by differential centrifugation to generate partially clarified medium (PCM) which contains all EV subtypes and was analyzed by nanoparticle tracking analysis (NTA). Top, a representative NTA plot of EVs shed from U87 cells with or without PTEN expression. The plot indicates the diameter and abundance of EVs within the PCM relative to cell number. Bottom, the total area under the curve from three different NTA experiments was used to calculate total EVs shed per cell. Two-way ANOVA using Tukey’s multiple comparisons test, ** p-value 0.0032. C) In a similar manner, two unique LN229 clones with confirmed CRISPR-mediated KO of PTEN (K1, and K2) and two control scrambled CRISPR treated LN229 clones (C1, and C2) were used to compare PI3K signaling and EV production. As before, cells were cultured in full serum media and washed three times with base media or PBS and then cultured in serum-free media for 12-15 hours. Cells were lysed and analyzed by immunoblot to compare PTEN, total AKT, phosphorylated AKT (T308, S473), and actin as the loading control. D) NTA was performed on the PCM of these cells from four different experiments to evaluate total EVs produced per cell. Two-way ANOVA using Tukey’s multiple comparisons test, *** p-value 0.0007.

In a similar manner, we next analyzed the ability of LN229 cells to produce EVs after knocking out *PTEN*. Isogenic LN229 cells were subjected to CRISPR-mediated PTEN KO, and the selected clones showed significant increases in p-AKT (Figure 2C) as well as a 40% increase in the number of EVs shed from the cells (Figure 2D). Comparable results were observed when we transiently knocked down (KD) PTEN in LN229 cells (Figures S2B-D). Together, these data suggest that the regulation of PI3K by PTEN is likely the predominant cause of reduced EV biogenesis.

### PTEN/PI3K signaling regulates PD-L1 and other EV associated cargo in glioma cell lines

As described earlier, treatment of U87, U251, and LN229 cells with IFN-γ led to an increase in PD-L1 at the cell surface. However, when these cells lacked PTEN, the levels of PD-L1 on their PM were elevated over their PTEN expressing counterparts (Figure 1C). As a result, we examined whether this would result in a corresponding increase of PD-L1 in their EVs. U87 cells with or without PTEN were treated with IFN-γ or PBS as a control for 24 hours. The cells were washed with serum-free media and cultured for 12-15 hours followed by generating PCM for each condition. To determine whether any changes in EV cargo were specific to a distinct subtype of EV, the PCM for each treatment condition was subjected to centrifugation at 15,000 x g to obtain large EVs (microvesicles) or what is referred to as the “P15” fraction. The resulting supernatant was then subjected to a subsequent ultracentrifugation step of 120,000 x g to obtain small EVs (exosomes) or the “P120” fraction (Figure S2A). Immunoblot analysis was performed on both whole cell lysates (WCL) and their respective P15 and P120 EV fractions to compare protein cargo. Because we had observed a reduction in EV biogenesis when manipulating PI3K signaling, we analyzed equal protein content for each EV subtype in our immunoblot experiments. As was previously the case, U87 cells exhibited an upregulation in PD-L1 expression in response to IFN-γ treatment in a PTEN-independent manner (Figure 3A). Interestingly, however, this increase in total PD-L1 levels only translated in an increase in U87 derived EVs if the cells lacked PTEN expression, this was more pronounced in the P120 exosome fraction (Figure 3B). To establish whether these effects were specific to PD-L1, we probed and blotted for other proteins such as the EGFR^51^ and Ras^52^ that have been previously reported to be included as cargo of U87-derived EVs. In fact, both signaling proteins exhibited a similar enrichment in the P120 EV fraction in the absence of PTEN (Figure 3B).

**Figure 3.**
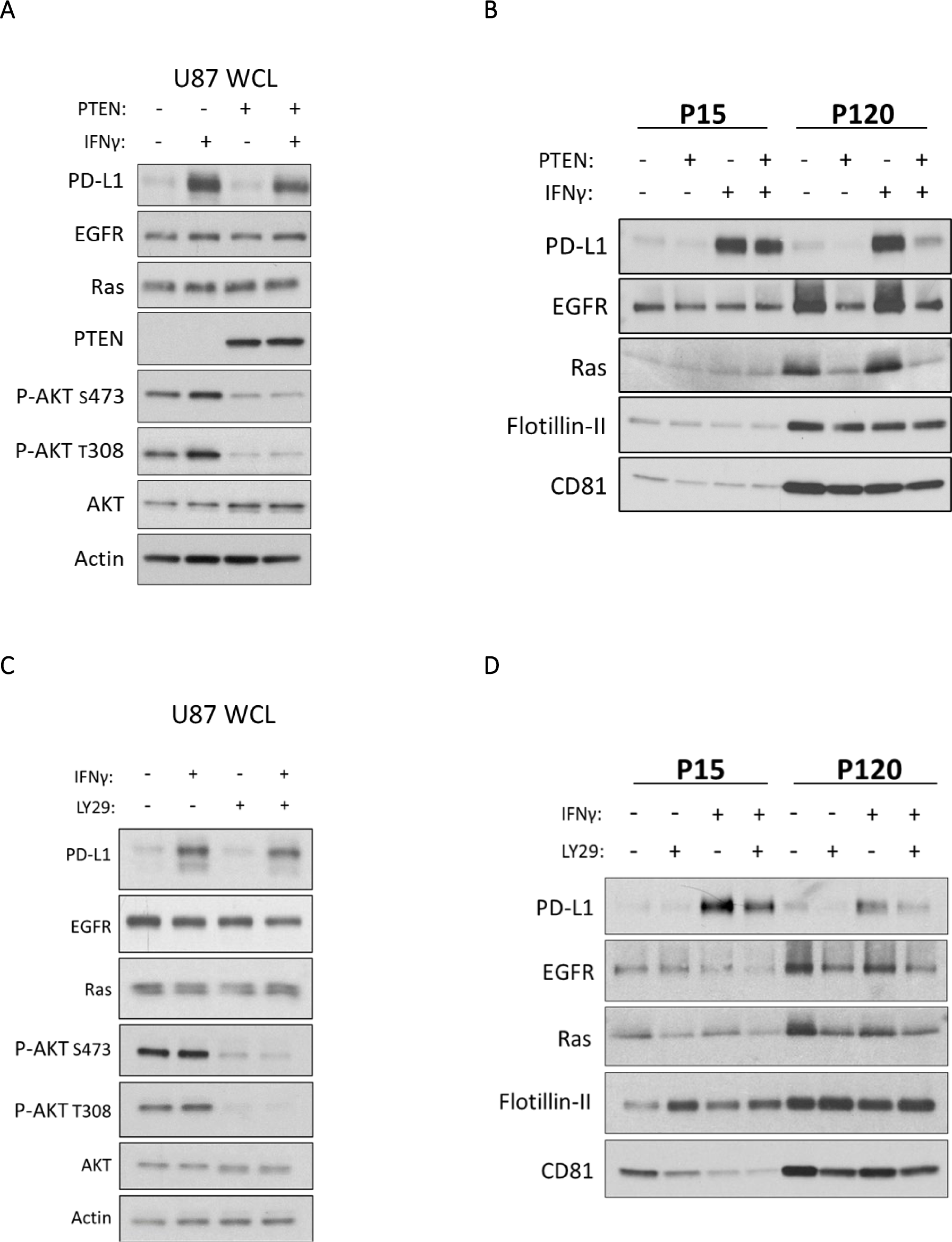
Inhibiting PI3K signaling negatively regulates PD-L1 enrichment in EVs shed from U87 cells. A) U87 cells, with or without PTEN, were treated with IFN-γ (50 ng/mL) or PBS as the vehicle control. Twenty-four hours later, cells were washed three times and cultured in serum free media for 12-15 hours. Whole cell lysates (WCL) were prepared and analyzed by immunoblot to evaluate expression of PTEN, AKT, phosphorylated AKT (T308, S473), and EV associated cargo of interest, including PD-L1, EGFR, and Ras. Actin served as a loading control. B) The PCM of each treatment group was subjected to centrifugation at 15,000 x g for 1 hour to collect the P15 or microvesicle fraction. The supernatant from this initial spin was collected and centrifuged for an additional 4 hours at 120,000 x g to collect the P120 or exosome fraction. The resulting immunoblot represents equal protein loading within each EV subtype. Each EV subtype was loaded to represent their relative abundance. PD-L1, EGFR, and Ras were probed as EV cargo of interest. CD81 was used as an exosome (P120) marker and Flotillin-II served as a loading control. C) Parental (PTEN deficient) U87 cells were treated with IFN-γ (50 ng/mL) or PBS for 23 hours. Cells were then pre-treated with the PI3K inhibitor LY294002 (20 μg/mL) for 1 hour, washed three times and incubated in serum-free medium supplemented with LY294002 (30 μg/mL) for 12-15 hours. Whole cell lysates (WCL) were subjected to immunoblot analysis to compare PD-L1, EGFR, Ras, AKT, and phosphorylated AKT (T308, S473), with actin as a loading control. D) Equal protein concentrations for both EV fractions were subjected to immunoblot analysis to compare PD-L1, EGFR, and Ras cargo. CD81 was used for confirming the isolation of small P120 EVs (exosomes) with Flotillin-II serving as a loading control.

No changes in CD81 and Flotillin-II were observed, suggesting that some specificity of cargo sorting existed. To establish whether these changes were occurring in a PI3K-dependent manner, we repeated these experiments with the PI3K inhibitor LY294002. As expected, immunoblot analysis confirmed that cells treated with LY294002 had significantly reduced p-AKT and cells treated with IFN-γ exhibited an upregulation of PD-L1 expression (Figure 3C). Analysis of the P120 EV fraction showed reduced quantities of PD-L1, EGFR, and Ras, while leaving Flotillin-II and CD81 unaffected (Figure 3D), similar to the results obtained upon ectopic PTEN expression.

To determine whether these results extended beyond U87 cells, we examined the effects of PTEN expression on PD-L1 associated cargo from our isogenic U251 and LN229 cell lines. In U251 cells, immunoblot analysis confirmed the PTEN-dependent reduction in p-AKT and the PTEN-independent upregulation of PD-L1 expression in both IFN-γ treated and untreated conditions (Figure 4A). As was observed for U87 cells, PTEN expression led to a marked reduction in PD-L1 and Ras cargo in the P120 EV fraction (Figure 4B). Reciprocally in LN229 cells, CRISPR-mediated PTEN KO resulted in elevated p-AKT (Figure 4C) and a corresponding enrichment of PD-L1 and EGFR in their derived EVs (Figure 4D). Collectively, these data suggest that PTEN loss leads to the enrichment of PD-L1 and other cargo within the P120 EV fraction in a PI3K-dependent manner. Indeed, when we expressed the constitutively active and oncogenic form of the EGFR (EGFRvIII) in U87 cells we saw an increase in PI3K/AKT activation (Figure S3A), a notable increase in PD-L1 cargo within the P120 EV fraction (Figure S3B) and in EV biogenesis (Figure S3C). The fact that we obtained highly similar results across multiple cell lines using both pharmacological and genetic approaches suggests that a conserved PTEN/PI3K dependent process broadly regulates both EV biogenesis and the loading of their associated cargo. Importantly, because the loss of PTEN leads to an increase in the numbers of EVs produced and an enrichment in the PD-L1 cargo associated with the vesicles, we hypothesized that these cells would inhibit TCR signaling to a greater extent relative to their PTEN expressing counterparts.

**Figure 4.**
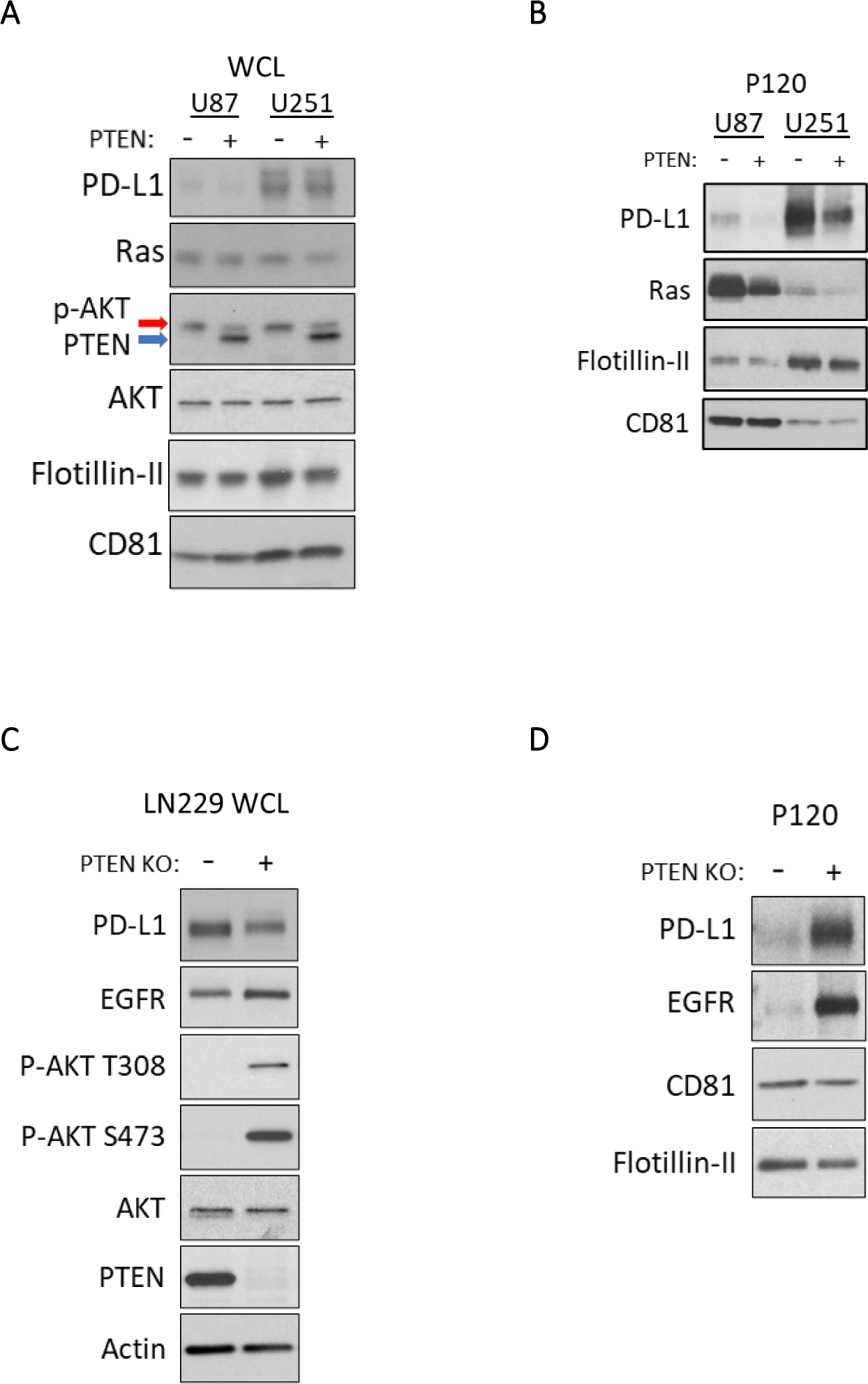
Altered PTEN expression in LN229 and U251 cells impacts EV associated PD-L1 cargo. A) Immunoblot analyses of U87 and U251 were carried out on WCLs to evaluate PTEN expression, p-AKT levels, and EV related cargo of interest including PD-L1, Ras, Flotillin-II, and CD81. B) The P120 EV fraction from these cells was collected as before, lysed, and analyzed by immunoblot analysis. Each condition had equal protein loaded to evaluate impact of PTEN expression on the enrichment of PD-L1, and Ras. Flotillin-II, and CD81 were used as EV and loading markers. C) LN229 PTEN KO clone 1 (K1) or LN229 CRISPR control clone 1 (C1) were treated with IFN-γ (50 ng/mL) for 24 hours and prepared for EV collection as above. Immunoblot analysis of WCL was performed for PD-L1, EGFR, PTEN, AKT, and p-AKT (S473, T308), with actin serving as the loading control. D) The P120 EV fraction was subjected to immunoblot analysis to compare PD-L1, EGFR, CD81 and Flotillin-II levels. All lysates were loaded using equal protein concentration.

### EVs shed from PTEN-null cells have an increased capability to inhibit TCR signaling

The activation of PD-1 by PD-L1 has been shown to recruit phosphatases that attenuate TCR signaling which can ultimately impair CD8^+^ T cell function^53^. Because we observed that cells lacking PTEN gave rise both to greater numbers of EVs, and an increase in the amount of PD-L1 that they contain, we hypothesized that EVs shed by these cells would be able to suppress TCR signaling to a greater extent than their PTEN expressing counterparts. To answer this question, we turned to the Jurkat lymphoma cell line as a well-established model system for studying TCR signaling.^54^ In order to quantify TCR activity, we used a luciferase Jurkat reporter cell line that is driven by the transcription factor nuclear factor of activated T-cells (NFAT). TCR-mediated activation of NFAT allows for the transcription of several T cell effector cytokines such as IL-2, IL-4, IL-10, and IFN-γ, and thus functions as a reliable proxy for TCR activity.^55^ Importantly, because Jurkat cells do not endogenously express the PD-1 receptor, we transduced the reporter Jurkat cells with PD-1 and used fluorescent activated cell sorting (FACS) to retain the highest expressors of PD-1, hereafter referred to as Jurkat^PD-1^ cells (Figures S4A and S4B). Finally, the function of the luciferase reporter system in the Jurkat^PD-1^ cells was confirmed by using a stimulating αCD3 (OKT3) monoclonal antibody with endpoints at 3-, 6-, and 20-hours post-treatment (Figure S4C). From these data, we calculated the EC50 for αCD3 to be 50 ng/mL which was used alongside the 6-hour time point for the remainder of our experiments.

We then examined whether glioma derived EVs were capable of inhibiting Jurkat^PD-1^ TCR activity by first utilizing the U251 cell line due to its robust expression of PD-L1 within their EVs relative to U87 cells (Figure 4B). Additionally, to confirm whether any observed TCR inhibition was occurring in a PD-1/PD-L1 specific manner, Jurkat^PD-1^ cells were pre-treated with an inhibitory αPD-1 antibody, or an isotype control antibody, 30 minutes prior to the simultaneous treatment with the stimulating αCD3 (OKT3) antibody and three increasing doses of EVs. Six hours post-treatment, cells were treated with luciferase substrate and luminosity readings were taken. Indeed, we observed that EVs from U251 cells caused a dose-dependent reduction in luminosity (TCR activity) which was rescued by the pre-treatment of Jurkat^PD-1^ cells with an inhibitory αPD-1 antibody (Figure 5A), indicating that these effects were specific to EV associated PD-L1.

**Figure 5.**
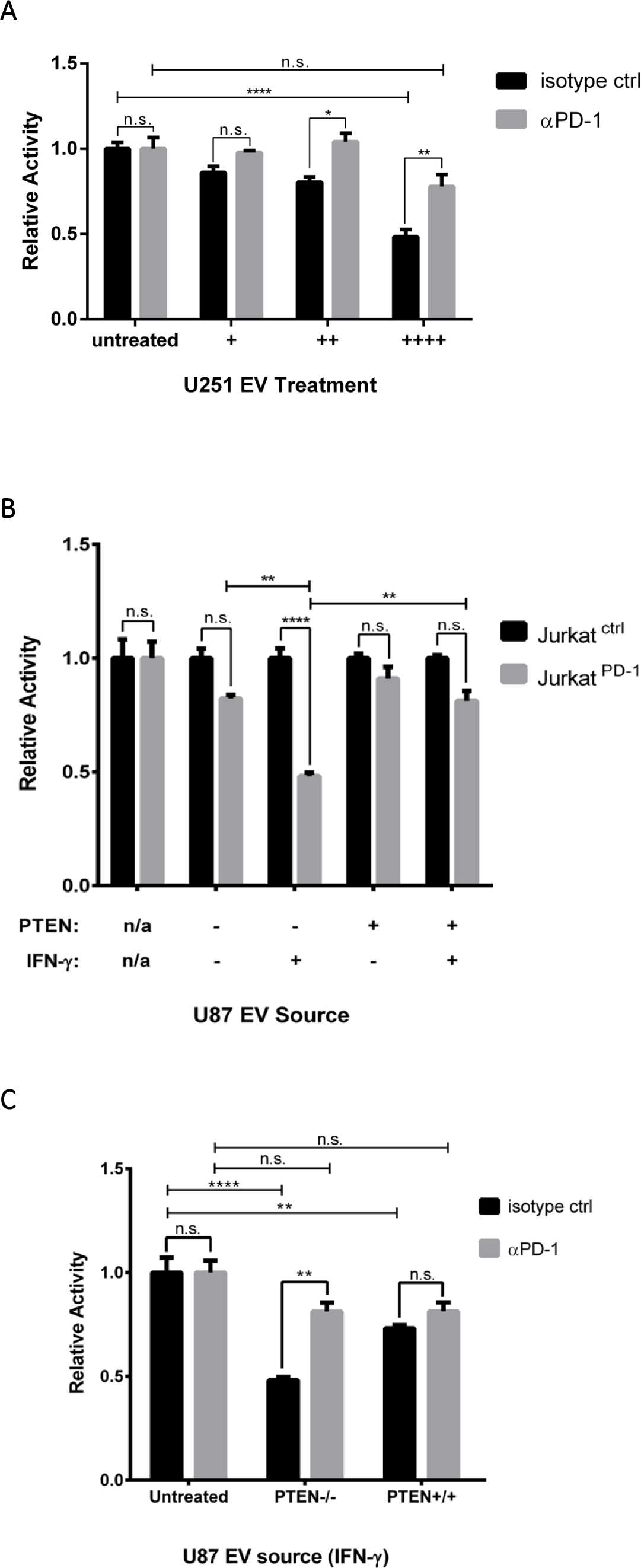
EVs from glioma cell lines abrogate TCR signaling in a dose dependent and PD-L1/PD-1 specific manner. A) Jurkat^PD-1^ cells were pre-treated with either an inhibitory αPD-1 or isotype control antibody 30 min before simultaneous treatment with EVs and activating αCD3 (OKT3) antibody at 50 ng/mL. EVs were sourced from U251 cells (no IFN-γ treatment) and split into three different doses. An EV treatment of (++++) corresponds to 4 x 10^8^ particles and was used to treat an individual well of 3 x 10^4^ Jurkat^PD-1^ cells, with (++) and (+) EV doses representing 1:2 and 1:4 dilutions, respectively, of the original dose. Luciferase values were normalized to isotype control Jurkat^PD-1^ cells for each condition. The Y-axis represents relative luciferase units (RLU). p-values: * 0.0398, **0.0035 **** < 0.0001. B) Jurkat^PD-1^ and Jurkat^Ctrl^ cells were simultaneously treated with OKT3 αCD3 antibody (50 ng/mL) and total EVs derived from U87 cells with or without PTEN expression and with or without IFN-γ (50 ng/mL). EV doses were sourced from 1.25 x 10^6^ U87 cells, representing 3 x 10^9^ particles from PTEN positive cells and 5 x 10^9^ particles from parental PTEN naïve U87 cells. EVs were used to treat 3 x 10^4^ Jurkat^PD-1^ cells and Jurkat^Ctrl^ cells which were used to normalize luciferase activity. C) To determine if EV mediated inhibition was specific to PD-L1, Jurkat^PD-1^ cells were pre-treated with αPD-1 antibody or isotype control antibody for 30 min before introducing the same EV treatment as above. Luciferase values were normalized to isotype treated Jurkat^PD-1^ cells for each treatment condition. Each experiment was done in triplicate and repeated twice. Statistical analysis was done by two-way ANOVA with Tukey multiple comparison. p-value **** < 0.0001, ** < 0.0080.

We then examined whether the IFN-γ promoted upregulation of PD-L1 would influence the magnitude of EV-mediated TCR inhibition, and if this was affected by the glioma cell’s PTEN status. To help us determine the specificity of PD-1/PD-L1 TCR signal attenuation, we treated Jurkat^PD-1^ and PD-1 naïve Jurkat^Ctrl^ cells with a moderate (++) dose of EVs shed from either PTEN expressing or non-expressing U87 cells treated with or without IFN-γ (see materials and methods). Overall, EVs derived from U87 cells treated with IFN-γ had a greater capability of inhibiting TCR signaling relative to the EVs shed from PBS treated cells (Figure 5B). More specifically, EVs from U87 cells expressing PTEN and treated with IFN-γ were only able to limit TCR signaling by 20%, compared to the 10% inhibition observed for EVs shed from the PBS treated control group. In contrast, EVs from parental U87 cells (PTEN-null) treated with IFN-γ led to a 50% reduction in TCR signaling, whereas EVs from the same cells without IFN-γ treatment, only resulted in 15% signal attenuation. Thus, PTEN expression negatively impacts EV-mediated TCR inhibition, essentially eliminating any benefits conferred by IFN-γ. To further illustrate that these inhibitory effects were specific to PD-1/PD-L1 function, EVs isolated from IFN-γ treated U87 cells, with or without PTEN expression, were used to treat Jurkat^PD-1^ cells that were either pre-treated with an inhibitory αPD-1 antibody or an isotype control antibody. Again, EVs from U87 cells lacking PTEN expression were significantly better at inhibiting TCR signaling; however, these effects were reversed when Jurkat^PD-1^ cells were pre-treated with the αPD-1 antibody (Figure 5C). Taken together, these data indicate that EVs can indeed suppress TCR signaling in a dose-dependent and PD-L1-specific manner. Most importantly, these data show that glioma cell lines lacking PTEN expression are likely better suited to evade immune detection in response to IFN-γ and implicate EVs as potentially important contributors in the formation of adaptive immune resistance.

## Discussion

GBM patients have long been recognized as being systemically immunosuppressed, with reduced T cell counts, circulating dendritic cells and monocytes.^56,57^ Because immune evasion is a hallmark of tumor progression, this suggests that tumor suppressors and oncogenes must have pleotropic roles that go beyond conferring cell-intrinsic benefits such as promoting cell growth^58^. For GBM, it is becoming increasingly clear that certain molecular and genomic characteristics have important yet undefined immuno-regulatory functions.^59,60^ In particular, the loss of PTEN has garnered increased clinical significance due to its prevalence in many cancer subtypes, its contributions toward an immunosuppressive TIME, and its recent appreciation for being a prognostic marker for patient response toward immune checkpoint inhibition^61^. However, the cell and molecular determinants by which loss of PTEN leads to these immunological phenomena are not well defined and thus an important area of focus. Studies have highlighted that loss of PTEN in glioma cells alters their secretome, leading to the production and secretion of distinct cytokines to drive the formation of critical feedback loops between tumor and myeloid cells.^30,31^ Additionally, PTEN loss has been shown to promote the evasion of T cell mediated killing due to an increase in the basal expression of PD-L1.^43^ However, cancer cells can also adaptively upregulate PD-L1 in response to T cell effector cytokines such as IFN-γ, further promoting their ability to evade immune detection and killing.^4^ Whether the signaling activities triggered by PTEN loss and IFN-γ could synergize to further promote the upregulation of PD-L1, was not clear and an initial motivator for this study.

Here, we use several cell lines and pharmacological inhibitors to investigate how these two signaling pathways independently and/or cooperatively impact PD-L1 expression. We found that in isogenic glioma cell lines, PTEN status has no impact on overall PD-L1 expression at the transcriptional level, which agrees with a study by Parsa and colleagues. ^43^ However, based on our Western blot analyses, we did not observe an increase in total cellular PD-L1 protein that the authors attributed to increased translation. It is possible that our data differs due to cell culture conditions as most of our experiments were conducted in the absence of serum which can negatively impact mTORC1 activity and thus translational machinery.^62^ Nonetheless, our study now highlights how IFN-γ affects PD-L1 expression in the context of PTEN status. We show that transcriptional upregulation of PD-L1 is dependent on JAK/STAT and not PTEN/PI3K signaling. Furthermore, our flow cytometry data reveal that PTEN expression negatively regulates plasma membrane-associated PD-L1 levels predominantly after exposure to IFN-γ. Overall, these findings agree with similar studies using colorectal and NSCLC cancer models.^45,47^ Importantly, because EVs either mature and bud off from the cell surface (i.e., large EVs, MVs, or P15), or are shed when MVBs fuse with the plasma membrane (small EVs, exosomes or P120)^63^, we hypothesized that our results were indicative of underlying changes in EV associated PD-L1 cargo and biogenesis.

While various lines of evidence have highlighted the role of EVs in PD-L1 mediated immune evasion,^40–42^ no study to date, to our knowledge, has evaluated whether PTEN/PI3K signaling can influence EV biogenesis and/or PD-L1 cargo. We now present evidence that links the loss of PTEN with enhanced EV biogenesis. These results were confirmed both by the ectopic expression of PTEN in U87 and U251 cells and by knock down and knock out experiments in LN229 cells. We also examined whether IFN-γ could alter EV biogenesis and if its ability to upregulate PD-L1 in different EV subtypes was dependent on PTEN expression. Although we did not observe any significant changes in EV biogenesis in response to IFN-γ treatment, we did see an enrichment in PD-L1 in both the larger P15 EVs (microvesicles) and smaller P120 EVs (exosomes) shed from glioma cells. Importantly, we show that PTEN expression in glioma cells significantly reduces the amount of PD-L1 present predominantly within the smaller P120 (exosome) EV fraction. Finally, because pharmacological inhibition of PI3K led to similar results as our genetic approach, we believe that these effects occur in a PI3K-dependent manner. We then developed a PD-1 expressing Jurkat NFAT reporter cell line to measure TCR signaling and found that EVs shed from glioma cells are capable of inhibiting TCR signaling in a dose-dependent and PD-L1-specific manner. Notably, cells treated with IFN-γ produced EVs with an enhanced capacity to inhibit TCR signaling. However, this improved immunosuppressive behavior was ameliorated if cells expressed PTEN. Overall, our work adds to a growing body of evidence implicating PTEN loss and/or hyperactive PI3K signaling with an altered cancer secretome associated with immuno-modulatory functions. Most importantly, our results highlight a previously unappreciated mechanism by which hyperactive PI3K in GBM, largely a result of PTEN loss, can regulate EV biogenesis and their associated cargos. Others have shown that EVs can also be enriched with other immunosuppressive proteins such as FasL, TRAIL, and galectin-9, which can promote T cell apoptosis, suppress TCR expression, and block cytotoxic functions of NK and CD8^+^ T cells.^64–66^ Thus, by enhancing EV biogenesis and influencing the recruitment of cargo into EVs, the loss of PTEN and/or hyperactivation of PI3K may thus lead to a considerable increase in the bioavailability of PD-L1 and other immuno-modulatory proteins in the extracellular milieu to ultimately contribute to a more immunosuppressive TIME.

Because PI3K inhibition yielded similar results as those obtained when PTEN was restored, it is possible that altered PIP dynamics, due to unregulated PI3K activity, is responsible for these EV associated changes. For example, the disruption of VPS34 (*PIK3C3)* activity, the sole lipid kinase that specifically generates PI(3)P, has been shown to lead to the release of atypical exosomes in the context of neurodegenerative disease.^67^ Similarly, the inhibition of the PIKfyve lipid kinase, which generates the endosome and MVB-specific PI(3,5)P2 from PI(3)P, has also been reported to give rise to increased exosome biogenesis.^68^ Given that PIP metabolism is subject to tight spatial/temporal regulation, wherein one PIP will act as a precursor for a lipid kinase or phosphatase, ^69^ disruptions in a single lipid kinase or phosphatase can have significant consequences on organelle identity, protein recruitment, endocytosis, endocytic trafficking, and the regulation of mitotic signaling events.^70,71^ Indeed, several studies have highlighted how PTEN and PI3K are dependent on distinct PIP species for their localization and function at the PM or within endosomal compartments.^72,73^ In those cases, PTEN and PI3K alter PIP species resulting in changes in endomembrane dynamics, altering membrane trafficking, protein recycling, endocytosis and endosomal maturation to regulate and propagate mitotic signaling.^50,74–76^ Thus, an increase in the levels of the PI3K generated phosphatidylinositol (3,4,5)-triphosphate (PIP3) at distinct stages of endocytic maturation and/or recycling, may in some way affect EV biogenesis and cargo sorting. Future studies will therefore be directed toward further elucidating the biochemical mechanisms by which altered PIP metabolism brought on by PTEN loss and/or oncogenic PI3K activity in discrete membrane compartments can impact endomembrane maturation, trafficking, and the recruitment of specific cargo into EVs.

## Materials and Methods

### Cells

U87MG, LN229, U251, and T98G cells were obtained from American Type Cell Culture (ATCC). Jurkat cells expressing the NFAT luciferase reporter were purchased from Promega (#J1601). Apart from Jurkat cells, all cell lines were maintained in Dulbecco’s Modified Eagle Medium (DMEM) supplemented with 10% fetal bovine serum (FBS). Jurkat cells were maintained in Roswell Park Memorial Institute 1640 (RPMI, Thermo #118750) medium supplemented with 10% fetal bovine serum (FBS, Thermo #16000044), 1 mM sodium pyruvate (Thermo #11360070) and 100 nM minimal non-essential amino acids (Thermo #11140050). Transduced semi-stable cells were cultured in their respective media supplemented with 250 ng/mL puromycin. Adherent cells were subcultured by washing once with 1x phosphate buffered saline (PBS) and dissociated by incubating cells with Hank’s balanced salt solution (HBSS) containing 0.25% Trypsin, 0.1% EDTA w/o Calcium, Magnesium and Sodium Bicarbonate (Corning #25053) for 3 minutes at 37°C. Complete media was used to quench trypsin activity followed by centrifugation of cells at 200 x g for 5 minutes. Jurkat cells were seeded at a density of 1 x 10^6^ cells per mL of media and subcultured every 3 days by collecting and pelleting floating cells at 400 x g for 5 minutes. In some cases, cells were treated with the JAK/STAT inhibitor Tofacitinib citrate (Cayman Chemical #11598), or with IFN-γ (Peprotech #300-02) or the PI3K inhibitor LY294002 (Cayman Chemical #70920) for the time-periods and concentrations indicated.

### RNA Isolation and Quantitative PCR Analysis

Total RNA from cells was isolated using Direct-zol RNA miniprep kits (Zymo Research, #R2051). RNA concentration was measured on a NanoDrop OneC (ThermoFisher) instrument. Total RNA (500 ng) was converted to cDNA using Superscript III Reverse Transcriptase (Invitrogen) and oligo dT20 primers. The cDNA was then used to determine the expression levels of the indicated transcripts using SYBR Green Supermix (Bio-Rad) in an Applied Biosystems ViiA 7 Real-Time PCR System and the ΔΔCT method. The following primers were used for the RT-qPCR analyses: PD-1

(CD274) Forward: GCTGCACTAATTGTCTATTGGGA

PD-1 (CD274) Reverse: AATTCGCTTGTAGTCGGCACC

β-actin Forward: CATGTACGTTGCTATCCAGGC

β-actin Reverse: CTCCTTAATGTCACGCACGAT

DNA constructs, cloning, subcloning and sequencing The pLJM1-empty (#91980) and PD-1-miSFIT-1x (#124679) vectors were purchased from Addgene. PTEN and PD-1 were subcloned into linearized pLJM1-empty vector using EcoR1 and NHE1 restriction enzymes (NEB) and Takara’s two-step In-Fusion® snap assembly (#638948) according to the manufacturer’s recommendations. *PD-1* was subcloned from the PD-1-miSFIT-1x construct using the forward primer: CGTCAGATCCGCTAGCCTGGAGACGCCATCCACG and reverse primer: TCGAGGTCGAGAATTCCGGAAGCTGCGCCTGTCATC. *PTEN* was cloned from cDNA generated from LN229 mRNA using forward (ATGACAGCCATCATCAAAGAGATCGTTA) and reverse (TCAGACTTTTGTAATTTGTGTATGCTGATCTTCATCAA) primers. The subcloned PTEN was then subjected to a second round of PCR using the forward primer (CGTCAGATCCGCTAGCCCACCATGGATTACAAGGATGAC) and the reverse primer (TCGAGGTCGAGAATTCTCAGACTTTTGTAATTTGTGTATGC). For all genes, the InFusion reaction mixture was used to transform Takara’s Stellar™ competent cells (#636766) using the heat shock method. For constructs needing higher DNA yields, Invitrogen One Shot™ Stbl3™ Chemically Competent *E*. *coli* cells were used (#C737303). To generate PTEN KO cells, two previously validated PTEN pLentiCRISPRv2 gRNAs^77^ were purchased through GenScript (ACCGCCAAATTTAATTGCAG and TTATCCAAACATTATTGCTA). Scramble lentiCRISPRv2 construct (Addgene, #169795) was used as a control sgRNA. Qiagen QIAprep Spin Miniprep Kit was used to isolate all plasmid DNA. DNA sequences were validated by Sanger sequencing at the Cornell Biotechnology Resource Center. All primers were purchased from Integrated DNA technologies (IDT)

For gene knockdown (KD) studies, MISSION® TRC shRNA Lentiviral Vectors (Sigma) were used.

LN229 cells were treated with five different PTEN shRNA constructs:

TRCN0000002746: CCGGCCACAGCTAGAACTTATCAAACTCGAGTTTGATAAGTTCTAGCTGTGGTTTTT

TRCN0000002748: CCGGCGTGCAGATAATGACAAGGAACTCGAGTTCCTTGTCATTATCTGCACGTTTTT

TRCN0000002749: CCGGCCACAAATGAAGGGATATAAACTCGAGTTTATATCCCTTCATTTGTGGTTTTT

TRCN0000230369: CCGGTATACCATCTCCAGCTATTTACTCGAGTAAATAGCTGGAGATGGTATATTTTTG

TRCN0000230370: CCGGCCACAAATGAAGGGATATAAACTCGAGTTTATATCCCTTCATTTGTGGTTTTTG

### Lentivirus production and transduction

HEK-293T cells were transfected using the FuGENE® 6 Transfection Reagent (Promega) following the manufacturer’s recommendations. Briefly, 33 μL of FuGENE® 6 solution was added to 570 μL of Opti-MEM™ reduced serum medium and incubated for 5 min. DR8.2 (1 µg) packaging construct (Addgene #8455) and the PMD2.G (5 μg) envelope construct (Addgene # 12259) were added to the solution, followed by the addition of 5 μg of either pLJM1-Empty, pLJM1-PTEN, pLJM1-PD-1, PTEN pLentiCRISPRv2, or shRNA constructs, and allowed to incubate for an additional 15 min. The transfection mixture was then added dropwise to 5 x 10^6^ 293T cells in a 10 cm tissue culture treated plate. The supernatant was collected 24 and 48 hours after transfection. Each collection was subjected to centrifugation at 500 x g for 10 min and pooled together. Viral supernatant was used fresh when possible or frozen at −80°C until needed.

All cells were transduced with a 1:1 mixture of viral supernatant and complete media, containing Polybrene (8 μg/mL) and 1x pen/strep for 8-12 hours. Jurkat cells were transduced by spinfection. Briefly, a 1:1 mixture of viral supernatant and complete media were incubated with 6 x 10^6^ cells for 30 minutes at 37°C followed by centrifugation at 300 x g for 30 minutes at 37°C. All cells were allowed to recover in complete media for at least 48 hours before antibiotic selection. Semi-stable cell lines were generated by selecting cells with puromycin (1-2 μg/mL) for 5 days, with media being replaced every 24 hours. On the sixth day, cells were maintained at 250 ng/mL puromycin. Cells treated with shRNA lentivirus were not selected using puromycin. Instead, following a 48-hour recovery period, cells were immediately used for their appropriate experiments such as EV collection and/or immunoblot analysis.

For genetic knockouts using CRISPR, LN229 and T98G cells lines were transduced as above with both PTEN pLentiCRISPRv2 gRNAs or with scramble lentiCRISPRv2 and treated exactly as described for adherent cells. After antibiotic selection, cells were sub-cultured at a density of 1.5 cells/well in a 96 well plate in the presence of 250 ng/ml puromycin to select for individual clones. Media was replaced every 3-4 days for 2 weeks. At the end of week 2, wells with single colony growth were sub-cultured and expanded. Knock outs of PTEN were confirmed by immunoblot analysis.

### Extracellular vesicle collection

For immunoblot analysis, 1.5 x 10^6^ semi-stable cells were seeded in three 15 cm tissue-culture treated plates (Corning) for each treatment condition used. The following day, cells were treated with IFN-γ (50 ng/ml) or PBS as a control. After 24 hours, cells were washed 3X with PBS or base media and cultured overnight for 12-15 hours. For each treatment group, three dishes were counted using a Bio-Rad TC20 cell counter to normalize EVs shed from cells by NTA, or by comparing protein changes through immunoblot analysis. EVs were collected by differential centrifugation (Figure S2A). The conditioned media was centrifuged at 300 x g for 5 minutes, transferred to a new tube, and subjected to a second spin at 500 x g for 10 min. This partially clarified media (PCM) was then transferred into 70 mL Beckman polycarbonate bottles (#355655) and subjected to ultracentrifugation at 15,000 x g (P15) for 1 hour using a Beckman type 45ti rotor. The supernatant was carefully removed and transferred to a new set of 70 mL polycarbonate bottles and centrifuged at 120,000 x g for 4 hours (P120). The initial P15 pellet was resuspended and washed in PBS and subjected to another spin in a Beckman TLA 100.4 rotor at 15,000 x g (19,000 RPM). The washed P15 pellet (microvesicles) was then resuspended in either RPMI for biological experiments or lysed for immunoblot analysis. The P120 pellet was also resuspended and washed in PBS and subsequently centrifuged in a TLA 100.4 rotor at 120,000 x g (54,000 RPM). The P120 pellet (exosomes) was either lysed or resuspended in RPMI for intended experimental analysis. In a similar manner, EVs for studies examining their effects on Jurkat cells were isolated by first seeding 5 x 10^5^ U87 or U251 cells in 10 cm dishes. EVs were collected from these cells by centrifuging PCM at 120,000 x g and washing the pellet in PBS. The resulting EV pellet was resuspended in RPMI and used as described below. All centrifugation steps were carried out in pre-chilled rotors at 4°C.

### Nanoparticle tracking analysis (NTA)

PCM was analyzed by nanoparticle tracking analysis on a Malvern Nanosight 3000 at the Cornell Nanofabrication Facility (CNF) with a camera level set to 12 and a threshold value of 7. Video was captured five times for 60 seconds for each sample. The total number of cells that shed EVs was used to normalize particle (EV) counts. Cells were acquired after collecting the conditioned media using the Trypsin subculturing technique described above and counted on a Bio-Rad TC20 automated cell counter.

### Immunoblotting

One confluent 15 cm dish of cells was lysed by using 1 mL of lysis buffer (25 mM Tris HCL pH 7.6, 100 mM NaCl, 1% Triton X-100, 1 mM DTT, 1 mM EDTA, 1 mM NaVO4, 1mM NaF, 1 mM β-glycerol phosphate, and 1 mg/mL each of aprotinin and leupeptin) followed by cell scraping to produce whole cell lysates (WCL). Two hundred μL of lysis buffer (without DTT) were used to lyse the total pool of EVs, or the individual P15 and P120 pellets. Protein concentration was analyzed using Bradford reagent (Bio-Rad, #5000006) with absorbance measured at 595 nm on a Beckman UV spectrophotometer. Unless otherwise noted, all lysates were subjected to SDS-PAGE on 4-20% TrisGlycine gels (Invitrogen). WCL and EV lysates were prepared with reducing or non-reducing 5X sample buffer, respectively, followed by boiling samples for 5 min. Lysates were loaded based on equal protein content (∼10-20 μg for WCL, and 2-5 μg for EVs). In one case, EV lysates were normalized by cell number and a proportional volume of lysate was loaded to illustrate the magnitude associated with the shift in EV biogenesis and cargo sorting. The separated proteins within the SDS gel were then transferred onto a 0.42 μm PVDF membrane. Membranes were blocked with 5% bovine serum albumin (Sigma) in TBST (19 mM Tris Base, 2.7 mM KCl, 137 mM NaCl, and 0.5 % Tween-20) for 1 hour and incubated with the indicated primary antibodies overnight at 4°C with gentle rocking. The following day, membranes were washed 3X with TBST for a total of 15 minutes and incubated with HRP conjugated secondary anti-mouse (7076S) or anti-rabbit (7074S) antibodies at a 1:5000 ratio (Cell Signaling Technology) for 1 hour at room temperature with gentle rocking. Membranes were then washed 3X with TBST for a total of 30 minutes, after which they were exposed to ECL reagent (Pierce) and imaged using photosensitive film. Except for β-actin (8H10D10), which was used a 1:5000 dilution, antibodies EGFR (D38B1), STAT3 (124H6), p-STAT3 S727 (6E4), PTEN (D4.3), total AKT (C67E7), p-AKT S473 (D9E), Ras (3965) and PD-L1 (E1L3N) were used at a 1:1000 dilution and purchased from Cell Signaling Technology. The CD81 antibody was used at 1:5000 dilution (R&D Systems, MAB4615).

### Flow cytometry & Fluorescence activated cell sorting (FACS)

To compare PD-L1 surface levels in isogenic glioma cell lines with or without PTEN expression, cells were seeded at 5 x 10^5^ cells in 10 cm tissue culture treated dishes (Corning). Cells were treated with IFN-γ (50 ng/mL) or PBS as the vehicle control for 24 hours. Cells were washed once with PBS and incubated with 3 mL of TrypLE (Gibco) for 3-5 min or until cells lifted off from the plates with gentle tapping. Cells were then treated with 7 mL of ice cold PBS, collected into 15 mL conical tubes, and centrifuged at 200 x g for 5 min at 4°C. Cells were resuspended in ice cold FACS buffer (5% BSA, 2 mM EDTA, 1% FBS in PBS) supplemented with 1:100 PE conjugated PD-L1 (eBioscience, MIH1), or with PE conjugated Mouse IgG1 kappa Isotype Control (P3.6.2.8.1), and incubated on ice for 30 min. Cells were washed twice using ice cold PBS followed by centrifugation at 200 x g at 4°C and fixed with 4% paraformaldehyde in PBS for 15 min. Cells were then washed twice by the addition of copious amounts of ice cold PBS and centrifugating as before. Flow cytometry was performed on a ThermoFisher Attune NxT instrument.

Semi-stable NFAT luciferase expressing Jurkat cells (pLJM1-empty and pLJM1-PD-1) were subjected to fluorescent activated cell sorting (FACS) to sort cells for PD-1 expression. Cells were pelleted at 400 x g for 5 min, resuspended in ice cold FACS buffer containing a 1:100 dilution of FITC conjugated PD-1 antibody (e-bioscience, MIH4), 1:500 fixable cell viability dye (e-bioscience, L34963), and incubated on ice for 30 minutes. Cells were washed by adding excess amounts of ice-cold PBS followed by centrifugation at 400 x g for 5 minutes. This was repeated and the resulting cell pellet was then resuspended in FACS buffer. Stained cells were sorted on a Sony MA900 with a 100 μm nozzle. Our gating strategy used empty vector Jurkat cells as a negative control for PD-1. To gate live/dead cells, a heated sample of Jurkat cells (1 h at 65°C) was used as a positive control. Jurkat cells were sorted into RPMI containing 50% FBS and 1x Pen/Strep. Sorting was stopped once 500,000 events was reached.

EV treatment of Jurkat reporter cell lines to measure TCR signaling FACS sorted Jurkat cells expressing the NFAT luciferase reporter, with or without PD-1 expression (Jurkat^PD-1^ and Jurkat^Ctrl^, respectively), were used to measure the impact of EV associated PD-L1 on T cell receptor (TCR) signaling. The resulting EV pellet for each experiment was resuspended in serum free RPMI. Three different EV doses were used to treat Jurkat cells in triplicate (3 x 10^4^ cells/well). For U251 cells, an EV treatment of (++++) corresponds to the total vesicles that originated from one confluent 10 cm dish of cells, or ∼3.8 x 10^6^ cells and 2.5 x 10^9^ EV particles being distributed across three triplicate wells or 9 x 10^4^ Jurkat cells. The two other EV doses, (++) and (+), were serially diluted with serum free RPMI at a 1:2 and 1:4 ratio, respectively. For U87 cells, a dose of (++++) corresponds to a confluent 10 cm dish, or roughly 2.5 x 10^6^ cells, and translates to about 6.4 x 10^9^ EV particles sourced from PTEN-positive cells and 1.05 x 10^10^ particles from parental PTEN-negative U87 cells. The two other EV doses, (++) and (+), were serially diluted with serum free RPMI at a 1:2 and 1:4 ratio, respectively.

All treatments conditions were prepared in advance in an ultra-low attachment 96-well plate (Corning, #3474) to allow the use of a multi-channel pipette. For the Jurkat luciferase experiments, only the inner 60 wells of a white 96-well clear bottom plate (Corning, #3610) were used. Positive control wells contained 25 μL of serum-free RPMI supplemented with 150 ng/mL of αCD3 antibody (eBioscience, OKT3). Baseline luciferase activity was measured with wells containing only RPMI and isotype control antibody. Experimental wells had 25 μL EVs (suspended in serum-free RPMI containing 150ng/mL of αCD3), sourced from either U87 or U251 cells. The plate was then placed in a 37°C incubator for 30 minutes prior to the addition of Jurkat cells. Following this step, Jurkat cells were seeded at a concentration of 6 x 10^5^ cells/mL or 30,000 cells per well at 50 μL/well. As such, all treatment groups had a final volume of 75 μL and a final αCD3 antibody concentration within the wells of 50 ng/mL. In some instances, Jurkat^PD-1^ cells were pre-treated with a functional grade anti-PD-1 monoclonal antibody (eBioscience, #J116), to mimic PD-1 checkpoint blockade, or with a mouse IgG1 kappa isotype control antibody for 30 minutes at room temperature. Jurkat cells were then incubated in a 37°C incubator for 6 hours, at which point, 70 μL of prepared luciferase substrate mix (Promega) was added to each well and incubated for 5 minutes at room temperature in the dark. Luminance values were read on a Tecan Spark Cyto plate reader.

### Statistical Analysis

Unless otherwise noted, all experiments were performed at least three times and averaged. Experiments where only two conditions existed, a two-tailed student T-test was performed. All other experiments with three or more conditions were analyzed by Two-way ANOVA with Tukey’s multiple comparison correction. Error bars represent the standard error margin (SEM). In all analyses, statistical significance was reached if the calculated p-value was > 0.05. Statistical analysis was completed on GraphPad Prism v7.

## Acknowledgements

We thank Elizabeth Burnett for secretarial assistance. This work was supported by a grant from the National Institute of Health (R01CA201402) to R.A.C. We graciously thank the Alfred P. Sloan Foundation for supporting J.C.S with a fellowship award. This work was performed in part at the Cornell NanoScale Facility, an NNCI member supported by NSF Grant NNCI-2025233.

## Author contributions

J.C.S., T.M.P., and D.A.Z. performed experiments presented in the manuscript. J.C.S., K.W., A.A. and R.A.C. were involved in writing the manuscript.

