## Supplemental Figures for "PTEN loss in glioma cell lines leads to increased extracellular vesicles biogenesis and PD-L1 cargo in a PI3K-dependent manner"

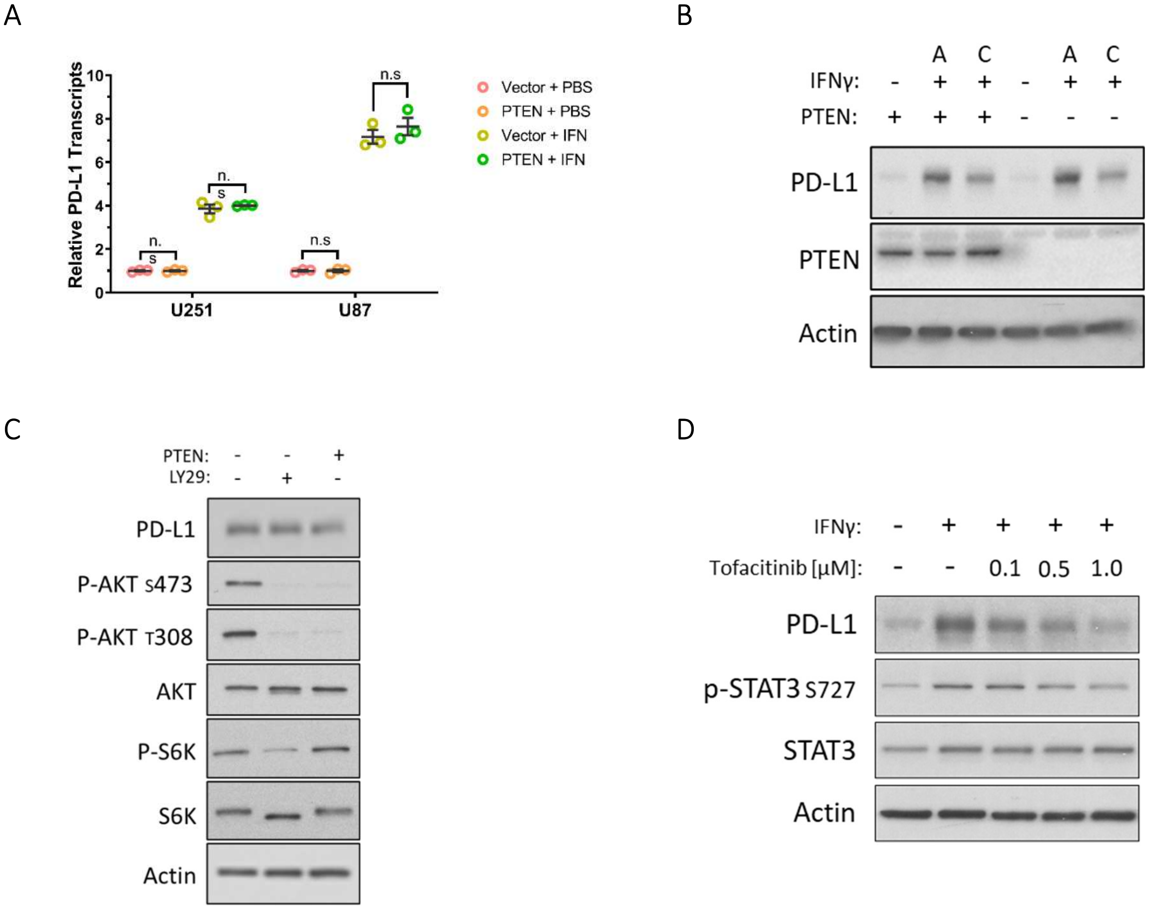


Supplemental Figure 1. IFN-γ upregulates PD-L1 in a PI3K/PTEN-independent manner

**A.)** Semi-stable isogenic U87 and U251 cells expressing either the control empty vector or PTEN vector, were treated once with IFN-γ (50 ng/mL) or PBS for 24 hours. RNA was isolated from each treatment group and subjected to reverse transcription (RT) PCR to generate cDNA and used to detect PD-L1 mRNA transcripts by qPCR. Within each cell type, Actin served as a housekeeping control to calculate the relative quantity for PD-L1 using the $2^{-\Delta\Delta Ct}$ method. **B.)** Semi-stable U87 cells expressing either the control empty vector or the PTEN vector, were either treated once with IFN-γ (50 ng/mL) for 15 hours, denoted with “A” for acute response, or cultured persistently with media supplemented with IFN-γ for five passages to represent a chronic state of inflammatory signaling and is denoted as “C.” Immunoblot analysis was performed to compare PD-L1 and PTEN expression with Actin serving as a loading control. **C.)** Two groups of U87 cells were transiently transduced with an empty vector (-) control and one group with the PTEN vector (+). Two days later, one empty vector control group was treated with the PI3K inhibitor LY294002 (LY29, 30 μg/mL), the other two groups were treated with DMSO as the vehicle control. All groups were concomitantly treated with IFN-γ (50 ng/mL). 24 hours after treatment, cells were lysed and immunoblot analysis was performed to compare PD-L1, PTEN, total AKT, AKT activity, (T308, S473), total S6K, S6K activity (T389) with Actin as the loading control. **D.)** In a similar fashion, U87 cells were concomitantly treated with the JAK/STAT inhibitor, Tofacitinib (CP-690550 citrate) or with DMSO (-) as the vehicle control, and IFN-γ (50 ng/mL). Immunoblot analysis was performed 16h post-treatment to compare PD-L1 expression and STAT3 activity (S727), with Actin serving as the loading control.

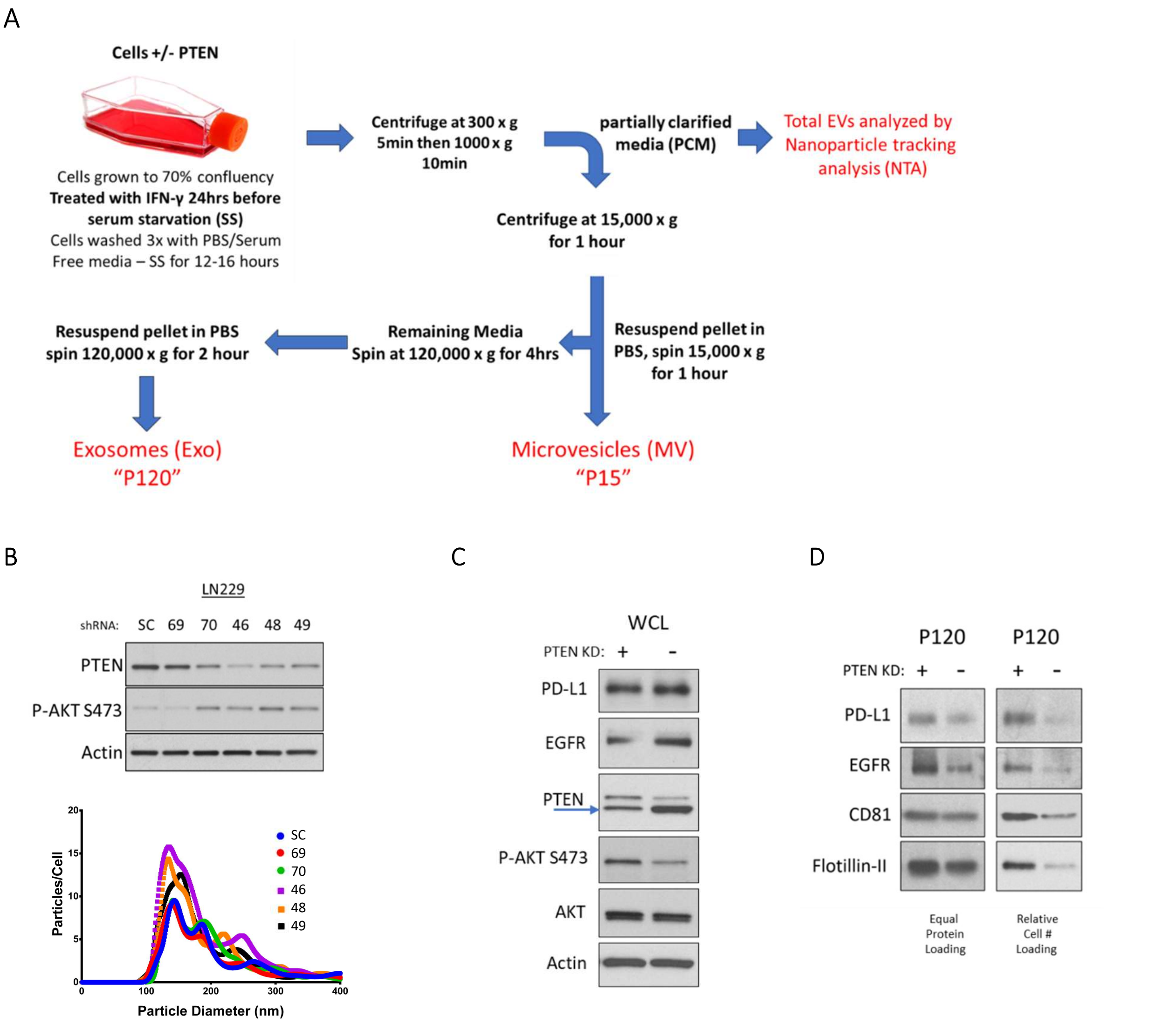


Supplemental Figure 2. KD of PTEN in LN229 cells leads to increased EV biogenesis and PD-L1 cargo

A.) Schematic of how cells were cultured and treated for EV collection. Partially clarified media (PCM) was used for EV analysis, and biological experiments, while the P120 and P15 fractions were utilized for immunoblot analysis. B.) LN229 cells were virally transduced with one scramble (SC) shRNA and five different shRNAs targeting PTEN. Each shRNA is represented by the last two digits of Sigma's MISSION® TRC shRNA numerical identifier as referenced in the materials and methods section. After 48 hours, cells were washed 3x with PBS or base media and then cultured for 12-15 hours in serum free media conditions. The supernatant was collected, and cells were washed, lysed, and subjected to immunoblot analysis to compare PTEN expression, total AKT, p-AKT (S473), and Actin as the loading control (top). The conditioned media from before was processed by differential centrifugation to produce PCM and subjected to NTA (bottom). C) LN229 cells were treated with PTEN shRNA (2746) or scramble shRNA through lentiviral transduction. 48 hours later, cells were treated with IFN-γ (50 ng/mL) for 24 hours and prepared for EV collection as before. Cells were lysed and subjected to immunoblot analysis using antibodies against PD-L1, EGFR, PTEN, AKT, p-AKT (S473) and Actin as a loading control D.) The P120 EV fraction (exosome) was subjected to immunoblot analysis for changes in PD-L1 and EGFR, while CD81 and Flotillin-II served as loading controls. (Left) blot loaded for equal protein content vs. (right) a blot loaded for relative EV output.

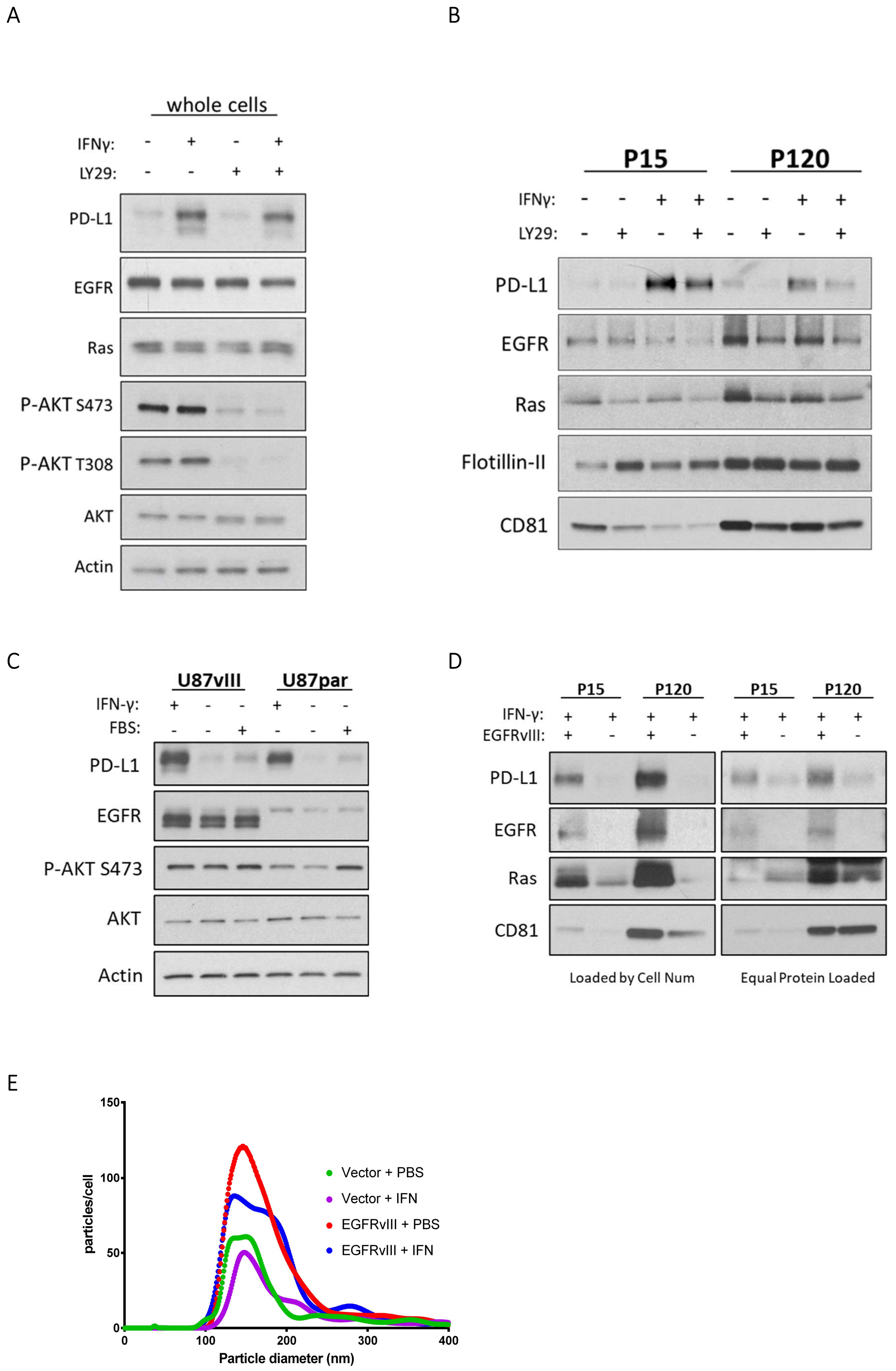

A
whole cells
IFNγ: - + - +
LY29: - - + +
PD-L1
EGFR
Ras
P-AKT S473
P-AKT T308
AKT
Actin
B
P15
P120
IFNγ: - - + + - - + +
LY29: - + - + - + - +
PD-L1
EGFR
Ras
Flotillin-II
CD81
C
U87vIII
U87par
IFN-γ: + - - + - -
FBS: - - + - - +
PD-L1
EGFR
P-AKT S473
AKT
Actin
D
P15
P120
P15
P120
IFN-γ: + + + + + + + +
EGFRvIII: + - + - + - + -
PD-L1
EGFR
Ras
CD81
Loaded by Cell Num
Equal Protein Loaded
E
particles/cell
150
100
50
0
0
100
200
300
400
Particle diameter (nm)
Vector + PBS
Vector + IFN
EGFRvIII + PBS
EGFRvIII + IFN

Supplementary Figure 3: PI3K activity in glioma cell lines leads to enriched PD-L1 associated EV cargo

A) Parental PTEN deficient U87 cells were treated with IFN-γ (50 ng/mL) for 23 hours. Cells were then pre-treated with the PI3K inhibitor LY294002 (20 μg/mL) for 1 hour. Cells were then washed three times and incubated in serum-free medium supplemented with LY294002 (30 μg/mL) for 12-15 hours. Whole cell lysates (WCL) were subjected to immunoblot analysis to compare PD-L1, EGFR, Ras, AKT, its active states (T308, S473), and Actin as a loading control. B) EVs were collected, washed, lysed, and subjected to immunoblot analysis to compare PD-L1, EGFR, and Ras cargo. CD81 was used for confirming EV segregation while Flotillin-II was used as a loading control. Equal protein was loaded within each EV subtype but loaded based on their relative abundance. C) Semi-stable U87 cells expressing EGFRvIII or empty vector parental cells were treated with IFN-γ (50 ng/mL) for 24 hours followed by 12-16 hours in serum free conditions. A separate group of cells was maintained complete media containing serum without IFN-γ. Cells were then subjected to immunoblot analysis to compare expression of PD-L1 EGFR, AKT, p-AKT (S473), and Actin, as the loading control. D) The PCM from the serum deprived cells was processed as before and analyzed by NTA. E) The PCM was further processed by differential centrifugation to isolate, the larger P15 EV fraction and smaller P120 EV fraction and analyzed by immunoblot analysis for PD-L1, EGFR, Ras and CD81 serving as loading marker. On the left, EV lysate was loaded based on cell count to help visualize the absolute shift in for our proteins of interest. On the right, a second SDS-PAGE was loaded based on equal protein content to visualize if cargo sorting shifted within the two EV fractions.

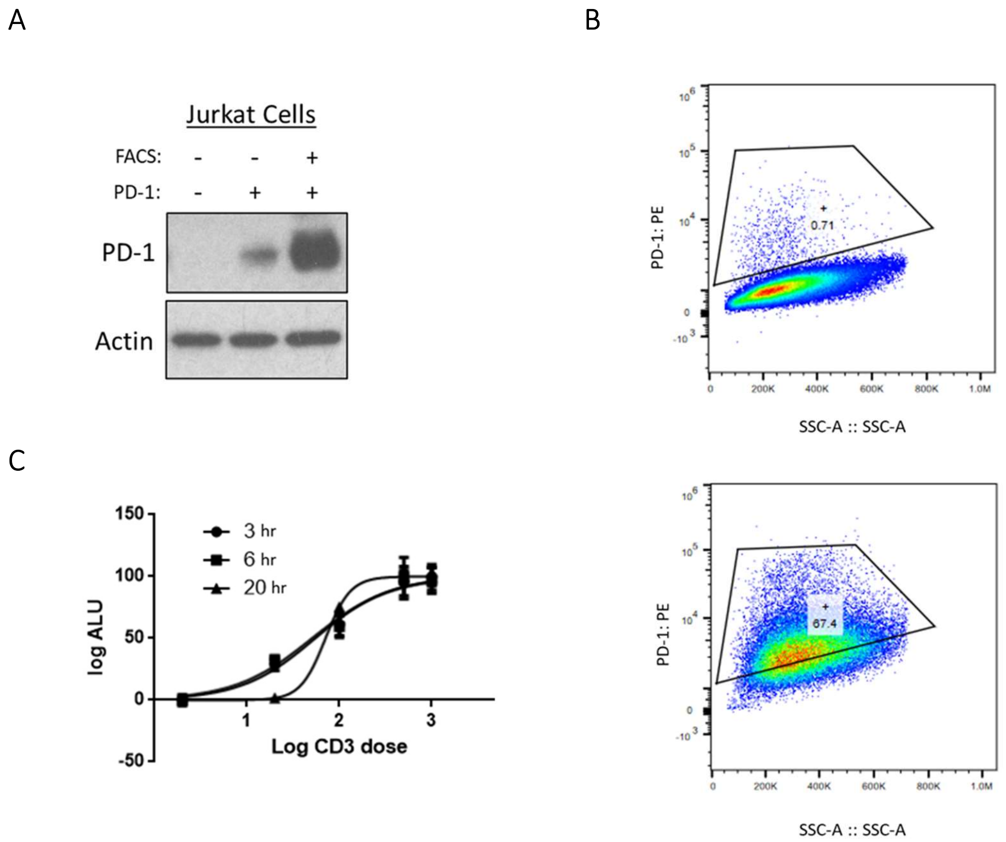


Supplemental Figure 4. Jurkat cells ectopically expressing an NFAT luciferase reporter and the PD-1 receptor respond to simulating αCD3 antibody

A) The Jurkat T lymphoma cell line was transduced with an NFAT luciferase reporter construct and a pLJM1-PD-1 or pLJM1-empty construct at a 1:1 ratio. Three days post-transduction, cells were analyzed by immunoblot for PD-1 expression and Actin as a loading control (lanes 1 & 2). PD-1 transduced Jurkat cells were enriched in B) and shown here. B) Transduced Jurkat cells were then sorted by FACS to enrich for high expressors for PD-1, and here after referred to as Jurkat$^{PD-1}$. Top, scatter plot showing pLJM1-empty vector transduced cells compared to the PD-1 sorted population, bottom. C) Jurkat$^{PD-1}$ cells were then treated with increasing concentrations of stimulating αCD3 antibody (OKT3) with three different end points of 3-, 6- and 20-hours post-treatment to determine an appropriate dose/time for our functional assays. From this, an EC50 for αCD3 antibody (50 ng/mL) was determined and used the 6-hour time point for the remaining of our assays.
